# Detection-Guided Beamforming for Efficient Bat Localisation

**DOI:** 10.64898/2026.09.06.749406

**Authors:** Roberto Alessandri, Lia Gilmour, Jin Jack Tan, Alejandro Osses, Dan Stowell

**Affiliations:** Tilburg School of Humanities and Digital Sciences, Tilburg University, Warandelaan 2, Tilburg, 5037 AB, The Netherlands; Bat Conservation Trust, 8 Battersea Park Road, London, SW8 4BG, United Kingdom; Sorama B.V., Achtseweg Zuid 153H, Eindhoven, 5651 GW, The Netherlands; Naturalis Biodiversity Center, Darwinweg 2, Leiden, 2333 CR, The Netherlands

**Keywords:** Acoustic beamforming, passive acoustic monitoring, bat echolocation, detection-guided processing, ultrasonic source localisation

## Abstract

Passive acoustic monitoring is widely used to study wildlife, but current approaches provide limited insight into the spatial behaviour of animals. In bat ecology, reconstructing flight trajectories is essential for studying habitat use, movement patterns, and interactions, yet it remains difficult to achieve under field conditions. Acoustic cameras offer a potential solution by enabling sound source localisation, but their practical application is limited by the computational cost of beamforming and by the non-stationary, broadband, and transient nature of echolocation calls. In particular, exhaustive beamforming over wide ultrasonic bandwidths and dense spatial grids becomes infeasible for continuous monitoring.

In this work, we propose a detection-guided beamforming framework for efficient localisation of free-flying bats. The method exploits the sparsity of echolocation signals by restricting beamforming to detector-identified time-frequency regions and combines this with physically consistent short-time analysis parameters and dense spatial sampling. The framework is evaluated on field recordings acquired with a Sorama CAM iV64s acoustic camera.

Results show that detection-guided processing reduces the number of beamformer evaluations by approximately 77×, corresponding to a reduction of about 98.7 % in computational effort, while preserving spatial resolution. At the same time, the proposed parameter configuration improves the stability and sharpness of reconstructed trajectories by avoiding artefacts associated with temporal averaging.

These findings demonstrate that high-resolution acoustic localisation can be achieved under realistic computational constraints, supporting the integration of acoustic cameras into ecological monitoring workflows. The proposed framework enables the extraction of spatial trajectories from passive acoustic monitoring data, facilitating spatially resolved analyses of bat behaviour in field conditions.

**Highlights:**

- Acoustic beamforming enables spatial analysis of bat behaviour but is computationally expensive.
- We propose a detection-guided beamforming framework using the BatDetect2 bat-call detector to exploit time–frequency sparsity.
- The method reduces beamforming cost by over an order of magnitude while improving trajectory stability.
- This enables near-real-time spatial localisation of free-flying bats in field conditions.

## 1. Introduction

Passive acoustic monitoring (PAM) is a key tool for quantifying biodiversity and understanding ecosystem dynamics at multiple spatial and temporal scales, i.e., by characterising how acoustic activity varies across locations and over time [1, 2]. With the increasing availability of passive sensors, PAM has evolved from small-scale studies to a central tool for large-scale ecological assessment, involving a pipeline that spans data acquisition, signal processing, and ecological inference [3]. As one of the most speciose and ecologically vital groups, bats represent approximately 50% of all terrestrial PAM research [4], serving as effective bioindicators for habitat quality in forest, riverine, and urban environments [5]. While standard approaches rely primarily on single-channel detection and classification to derive activity indices [6], acoustic cameras (dense microphone arrays) enable sound sources to be localised in space rather than only detected over time, supporting the analysis of animal movement, behaviour, and habitat use [7, 8, 9, 10]. Existing acoustic localisation systems range from distributed microphone networks based on time-difference-of-arrival to more compact arrays performing direction-of-arrival estimation [11], with acoustic cameras representing a high-resolution extension of the latter. Such arrays provide a non-intrusive alternative to physical tagging while enabling simultaneous tracking of multiple individuals to study social interactions or estimate species abundance [12].

In the context of bat ecology, such spatial information is essential for reconstructing flight trajectories, studying echolocation behaviour, and investigating habitat use, movement patterns, and species interactions relevant to ecological monitoring and conservation applications [12, 7]. However, despite this potential, the application of acoustic cameras to continuous monitoring remains limited in practice [13] due to substantial barriers in hardware cost, the complexity of sensor synchronisation, and the computational burden of processing large volumes of high-frequency data [12, 7], a challenge that becomes increasingly significant as PAM moves towards large-scale deployments requiring continuous data acquisition and analysis [3].

If microphone arrays are used, a core challenge arises from the characteristics of bat echolocation signals. Calls are short and highly non-stationary in both time and frequency [14, 8], and in addition, bats are also fast-moving sources [15]. This rapid motion introduces Doppler-based time warping and unequal phase shifts across microphones in arrays [8], leading to time-varying propagation effects that violate the quasistationarity assumption underlying frequency-domain beam-forming [16]. These factors impose strict constraints on short-term analysis parameters and spatial sampling requirements for accurate localisation, in order to avoid “ghost” sources and spatial smearing [12].

These physical constraints complicate the application of standard array-processing techniques in real-world conditions and limit the practical deployment of microphone arrays for continuous PAM. High-resolution beamforming over ultrasonic bandwidths further increases this challenge due to its computational cost. While recent advancements in deep learning have improved automated identification [17, 2], the labour-intensive steps of spatial position estimation have largely remained unautomated [12], and existing advanced methods often require significant technical expertise, limiting their accessibility for ecologists and practitioners without background in signal processing or machine learning [18]. Exhaustive processing scales with the joint time-frequency and space dimensionality, making continuous analysis impractical for long recordings. As a result, there is a need for approaches that reduce computational cost while preserving localisation accuracy.

In this work, we propose a detection-guided framework, implemented using BatDetect2, a spectrogram-based bat call detector, leveraging its time-frequency localisation and real-time inference capabilities [19, 20]. Detection outputs are used to constrain the time and frequency regions over which beamforming is performed, thereby reducing the number of required evaluations. This strategy is combined with a systematic analysis of short-time analysis parameters and spatial sampling to ensure that localisation remains physically consistent under the non-stationary conditions characteristic of bat echolocation. By decoupling localisation fidelity from computational cost, this framework addresses one of the aspects of the “nightmare” of protocol heterogeneity in large-scale PAM datasets [6] and supports the deployment of acoustic cameras for routine biodiversity monitoring [4, 5].

We evaluate the proposed framework using field recordings of free-flying bats. The results indicate substantial reductions in computational load while preserving spatial resolution under physically consistent parameter choices.

The remainder of this paper is structured as follows. Section 2 introduces the theoretical background underlying beamforming and its computational constraints. Section 3 describes the data acquisition and processing pipeline. Section 4 presents experimental results, followed by discussion and implications in Section 5. Conclusions are drawn in Section 6.

## 2. Theoretical Background

This section establishes the physical and computational principles underlying high-resolution ultrasonic beamforming of bat echolocation signals, focusing on array processing, time-frequency constraints, spatial sampling requirements, and computational scaling. An overview of the processing pipeline is shown in Fig. 1. The analysis in this work assumes far-field and free-field propagation conditions, under which delays depend only on the direction of arrival.

**Figure 1:**
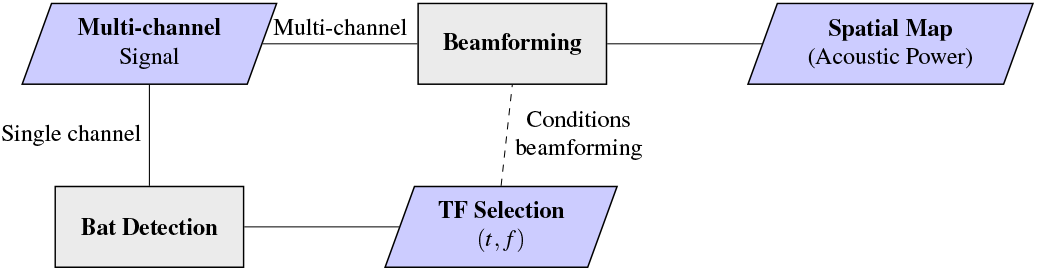
Detection-guided beamforming pipeline. A single channel is used for bat detection, generating time–frequency (TF) regions of interest. These regions condition the beamforming stage, which operates on the full multichannel signal and produces a spatial acoustic map.

### 2.1 Beamforming formulation

Frequency-domain delay-and-sum (DAS) beamforming is a widely used and physically interpretable baseline for acoustic imaging [16, 21, 22]. Under far-field assumptions, signals received at the array can be modelled using direction-dependent delays, and spatial localisation is obtained by evaluating a pseudospectrum over a predefined angular grid. When evaluated over a spatial grid, the peaks of this pseudospectrum correspond to candidate source directions. Additional details are provided in Appendix A.

For a recording segmented into *K* short-time frames, with *N*_*f*_ frequency bins per frame and a spatial grid of *N*_dir_ candidate source directions, exhaustive beamforming evaluates *P*(*θ, φ, ω*) for every frame, where (*θ, φ*) denote the candidate source direction and *ω* the frequency bin under analysis. The total number of beamformer evaluations therefore scales as *K*× *N*_*f*_ ×*N*_dir_, corresponding to all time-frequency-direction combinations to be processed.

This exhaustive evaluation is a key cause of the intrinsic computational cost of beamforming, which grows rapidly with increasing temporal, spectral, and spatial resolution.

### 2.2 Time–frequency constraints and motion effects

Frequency-domain beamforming assumes that source position and propagation geometry remain approximately constant over the duration of the analysis window [16]. If this assumption is violated by source motion, the phase relationships across microphones become time-varying, leading to a mismatch with the fixed steering vector. As a result, coherent summation is degraded, causing reduced mainlobe amplitude, energy spreading across directions, and loss of spatial localisation accuracy.

Bat echolocation calls are highly transient, with most UK species emitting frequency-modulated (FM) calls of duration 1− 5 ms, as illustrated in Fig. 2, and flying at 3 −5 m/s, with speeds up to 8≈ m/s [23, 24, 15]. These calls can range from below 25 to over 90 kHz, but usually occupy 40 −70 kHz. In addition to short call duration, echolocation call rates are high, typically ranging from approximately 10 to 40 calls per second. At the upper end of this range, successive calls may be separated by less than 25 ms, as shown in Fig. 2.

**Figure 2:**
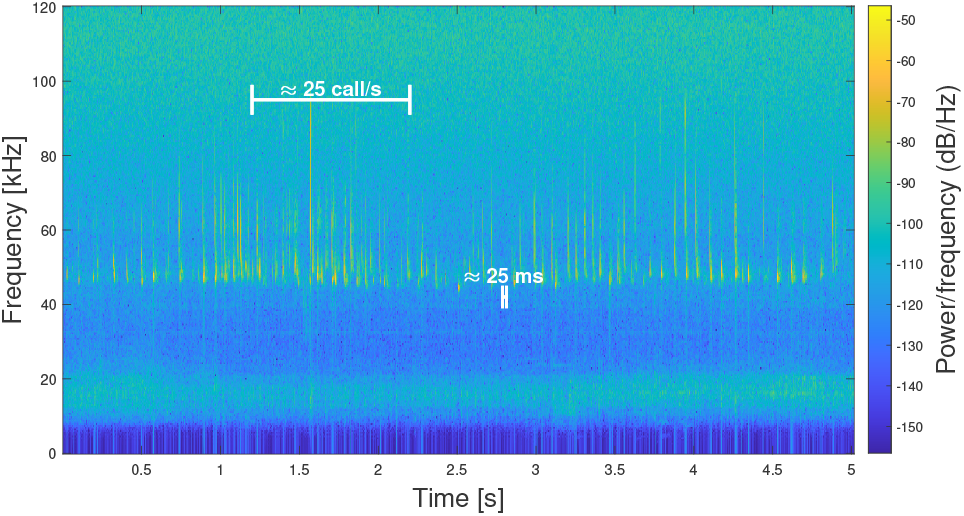
Example Spectrogram containing bat calls (high-passed at 10 kHz).

Large hop sizes risk missing entire calls or capturing them away from their peak energy. Consequently, overlap must be sufficiently high to ensure reliable temporal coverage.

Source motion constrains the maximum permissible window length. For a source at range *R*, moving with velocity *v* at incidence angle *Ψ*, the angular displacement during a window of length *L* samples is approximately

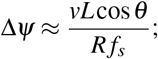

where *f*_*s*_ denotes the sampling frequency. When this angular displacement becomes comparable to the array beamwidth, localisation performance degrades due to spatial smearing. For a window duration of *T* = 17 ms and bat velocities at the upper end of the observed range, the resulting displacement is sufficient to produce noticeable localisation bias under typical recording conditions.

### 2.3 Spatial sampling and resolution

Beamforming produces a discrete spatial pseudospectrum (beamforming map) evaluated on an angular grid. If the grid spacing is too coarse, the source peak may fall between grid points, leading to underestimated levels and reduced localisation accuracy, as illustrated by the beamforming map examples in Fig. 3. These considerations show dense grids are not a design choice but a physical requirement, as insufficient sampling leads to spatial aliasing and loss of localisation accuracy [11].

**Figure 3:**
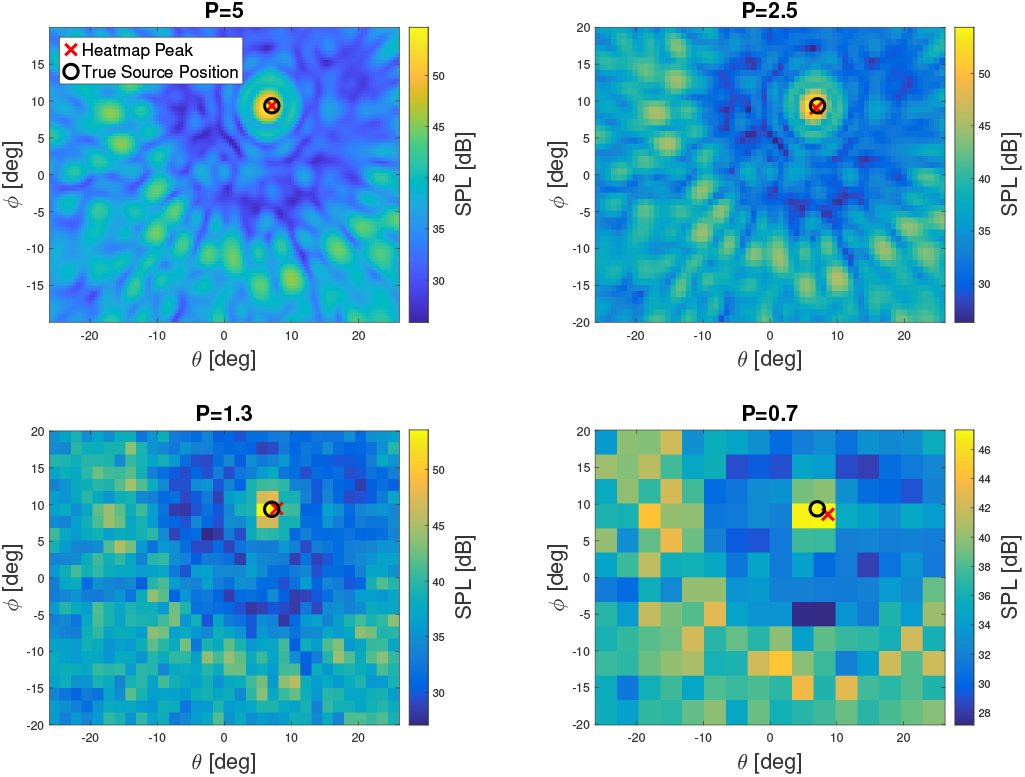
Beamforming pseudospectra for decreasing spatial oversampling factors *P*. Higher oversampling improves peak localisation and reduces discretisation artefacts.

For a planar array of effective aperture *D*, the angular resolution (half-power beamwidth) scales as

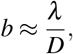

where *λ* is the acoustic wavelength [16]. At ultrasonic frequencies (e.g., 90 kHz, *λ* ≈3.8 mm) and *D* = 0.15 m, the beamwidth is approximately 1.46^°^ degrees. For reference, at a range of 10 m, this corresponds to a spatial resolution of approximately 25 cm, illustrating the need for dense angular sampling.

To adequately sample the beampattern, the angular grid spacing must satisfy 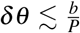, where *P* is an oversampling factor (typically 4 − 5) [25]. Over a total angular extent Ψ_Fov_,

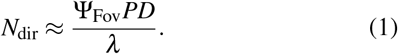

For example, assuming a field of view of Ψ_Fov_ = 120^°^, with *D* = 0.15 m, and *λ*≈ 0.0038 m, this yields approximately 412 grid points per angular dimension. Dense grids are therefore required for accurate peak localisation.

### 2.4 Computational scaling and sparsity

Frequency-domain delay-and-sum beamforming is computationally dominated by the spatial steering operation. For *K* time frames, *N*_*f*_ frequency bins, *N*_dir_ spatial points (total number of angular grid locations) and *M* microphones, the computational complexity scales as *O*(*KN*_*f*_ *N*_dir_*M*).

Given *α* the overlap, with 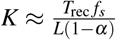, corresponding to the number of analysis frames obtained from a recording of duration *T*_*rec*_ with window length *L* and *N*_*f*_ ≈*L/*2, the combined time–frequency factor scales as

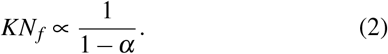

Consequently, short analysis windows required for quasistationarity do not inherently increase computational cost.

At ultrasonic frequencies, the dominant scaling is driven by *N*_dir_, as dense spatial grids are required to resolve narrow beams.

Memory requirements scale as *O*(*N*_*f*_ *N*_dir_*M*) ≈ *O*(*LN*_dir_*M*), so both spatial resolution and window length directly impact storage. For high-resolution grids and ultrasonic sampling rates, precomputed steering vectors can exceed the memory budget of embedded or edge platforms.

Taken together, these observations show that computational cost is fundamentally driven by the joint time-frequency-space dimensionality of the problem, motivating strategies that reduce this dimensionality without compromising spatial resolution.

### 2.5 Existing beamforming approaches and limitations

Beamforming approaches based on exhaustive evaluation of the search space include high-resolution methods [26, 27], which are effective but can face significant challenges due to intractable computational costs even in simpler contexts. In addition, frequency-domain Delay-and-Sum (DAS) beamforming provides a physically interpretable and widely used baseline [16, 22, 28], but when applied exhaustively over time, frequency, and dense spatial grids, its computational cost scales with the joint time-frequency-space dimensionality, leading to poor practical scalability. Post-hoc deconvolution techniques can improve spatial resolution but introduce substantial additional computational cost due to their iterative nature and their reliance on accurate steering vectors [25, 29, 30]. These methods additionally suffer from practical limitations, including sensitivity to steering mismatch and synchronisation errors under field conditions [7]. As a result, existing methods often face a trade-off between spatial resolution, computational tractability, and robustness.

An alternative strategy is to reduce the problem size before beamforming by exploiting the sparsity of acoustic scenes. Detection-guided localisation has been proposed as a means of restricting processing to time regions where relevant objects are present [31]. This redefines sound source localisation as a sparse optimisation problem for which recent data-driven approaches have attempted to mitigate its grid-related constraints using deep learning [32], yet require predefined knowledge of source counts. While object detection-guided beamforming has been previously proposed, its interaction with short-time analysis constraints and spatial sampling requirements in highly non-stationary ultrasonic regimes remains underexplored. For this approach to be efficient, the computational cost of the detection stage must remain small relative to that of the beamforming stage. This methodology aligns with current trends in ecoacoustics toward moving from purely descriptive studies to mechanistic, process-based analyses that can be applied routinely at scale, including applications in habitat monitoring, behavioural ecology, and conservation [1, 2]. Taken together, these limitations highlight the need for methods that reduce computational cost without compromising spatial resolution, motivating the detection-guided strategy developed in this work.

## 3. Materials and Methods

This section describes the data acquisition and processing pipeline used to implement the detection-guided localisation framework under realistic field conditions.

### 3.1 Recording sessions

Recordings were collected across multiple field sessions conducted at different locations and dates in London, UK, under varying environmental and acoustic conditions. Measurements were carried out in September 2025 in open environments, such as urban green spaces, selected based on expected bat activity, such as proximity to water bodies and open flight paths. More details in Table 1.

**Table 1:** Number of detections and unique species for each recording session.

| Session | Location | N. Detections | N. Species |
| --- | --- | --- | --- |
| 1 | Site A | 175 | 6 |
| 2 | Site A | 162 | 4 |
| 3 | Site B | 30 | 4 |
| 4 | Site C | 134 | 3 |
| 5 | Site D | 115 | 3 |

Recordings began approximately 20 minutes before sunset and lasted between 1 and 3 hours and consisted of multiple short recordings. In total, 5 recording sessions were processed, comprising 616 individual files. The number of bats ranged from single individuals to groups of more than 3 per recording. Environmental conditions, including temperature (12− 19^°^*C*), humidity (60 −95%), and wind levels, varied across sessions. Recording sessions generally avoided heavy rain; however, wind conditions still fluctuated, with maximum wind speeds reaching 34 km/h. Detected calls are consistent with common urban UK bat taxa, predominantly *Pipistrellus* species (*P. pipistrellus, P. pygmaeus, P. nathusii*), with occasional detections consistent with *Nyctalus* species. Species classification was not used in the subsequent analysis.

Signal quality varied across datasets, with mean Signal-to-Noise Ratio (SNR) values per session ranging from approximately 5 dB to 20 dB (Appendix B for details on the estimation procedure). This variability provides a representative dataset for evaluating localisation under realistic field conditions.

### 3.2 Acoustic camera and acquisition setup

The recordings were acquired using a Sorama CAM iV64s acoustic camera, comprising a 64-microphone planar sunflower (Vogel spiral) array with a circular aperture of 0.15 m, shown in Fig. 4. The sunflower geometry was used because it provides a near-Pareto-optimal trade-off between main-lobe width and sidelobe suppression while mitigating grating lobes [33, 34].

**Figure 4:**
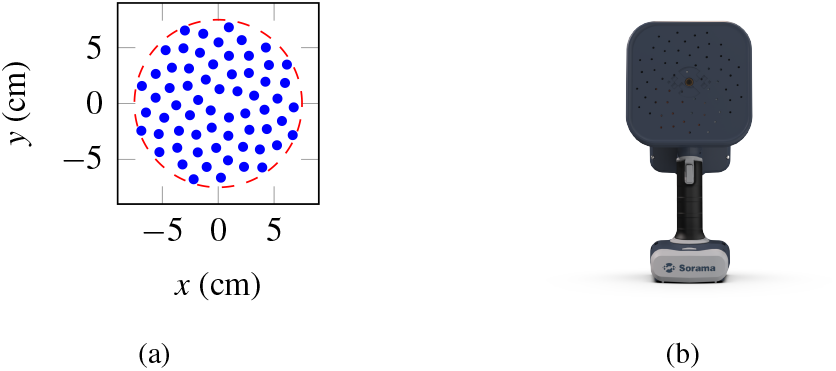
(a) Sunflower microphone array geometry used in the Sorama CAM iV64s. (b) Front view of the corresponding device.

All microphones recorded synchronously at 240 kHz (Nyquist frequency 120 kHz) for sessions 3, 4 and 5, sufficient to capture almost all frequency-modulated bat calls of UK species. For sessions 1 and 2, the sampling frequency was 120 kHz.

Recordings were conducted handheld, with the operator visually tracking bats using the device’s integrated camera, which has an approximate field of view (FoV) of ±20^°^ in azimuth and ±26^°^ in elevation.

Manual motion was kept smooth and slow such that the array can be treated as quasi-stationary within each analysis window.

### 3.3 Bat detection

Bat echolocation calls were detected using a single reference microphone channel, reducing computational cost while preserving reliable event timing for multichannel beamforming.

Detection was performed using BatDetect2 [19]. A high detection threshold (0.759) was adopted to prioritise precision over recall [20], resulting in sparse but reliable detections. Only detection outputs were used in the present work; species classification provided by BatDetect2 was utilised only for the example traces in Fig. 7.

To verify timing consistency, BatDetect2 was applied independently to each channel and detections were grouped using a tolerance window Δ*t*_*c*_. Consensus times were defined as median values within clusters.

### 3.4 Detection-guided beamforming

We propose a detection-guided beamforming framework in which beamforming is restricted to time-frequency regions identified by an automatic detector.

Let *ρ*_*t*_ denote the fraction of active time frames relative to the total number of frames, and *ρ*_*f*_ the fraction of retained frequency bins. Since bat echolocation calls occupy only a small portion of recordings in both time and frequency, these factors are typically *ρ*_*t*_ ≪1 and *ρ*_*f*_ ≪1.

Under this formulation, the effective number of beamformer evaluations is reduced from *O*(*KN*_*f*_ *N*_*dir*_) to

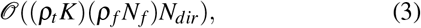

yielding a multiplicative reduction in computational cost.

The detection stage is implemented using BatDetect2, and the resulting time-frequency constraints are used to guide beamforming as illustrated in Fig. 1.

Spatial localisation was performed using delay-and-sum beamforming applied selectively in time and frequency based on detector output and compared with non-detection-guided beamforming.

Spatial pseudospectra were evaluated over grids of 36 ×48 points over a ±20^°^× ±26^°^ Fov. This configuration was selected to make runtime analysis computationally tractable while preserving the scaling behaviour of the beamforming pipeline.

### 3.5 Short-time analysis parameters

Signals were segmented using a short-time Fourier Transform with window lengths *L*∈ {256, 512, 1024, 2048, 4096} samples (1.07–17.07 ms at 240 kHz) and overlaps of 0% 25%, 50%, and 75%.

A configuration of *L* = 1024 and 75% overlap was selected. This corresponds to a window duration of 4.27 ms and frequency resolution of approximately 234 Hz, providing a balance between temporal resolution, frequency resolution, and quasi-stationarity.

### 3.6 Empirical assessment of temporal sampling

The selection of Short-Time-Analysis Parameters represents a trade-off between temporal resolution, signal support, and trajectory stability. While Sec. 2.2 establishes theoretical constraints based on quasi-stationarity and source motion, the practical impact of window length, overlap, and frame selection criteria was evaluated empirically. As no ground-truth trajectories are available for the recorded data, the objective is not to estimate localisation accuracy in absolute terms, but to compare physically plausible parameter configurations under identical recordings. The evaluation is therefore based on relative comparisons across parameter configurations, with consistency and physical plausibility used as primary indicators of performance. To provide an intuitive representation of the localisation output, Fig. 5 shows a two-dimensional spatial trajectory corresponding to a single recording. This representation illustrates the reconstructed flight path in Polar coordinates before analysing the individual Azimuthal (*φ*) and Elevation (*θ*) components as functions of time. In this and all following traces, the coordinates used for each location are the centroids of all the energy values between the maximum and −3dB in the map of the corresponding frame.

**Figure 5:**
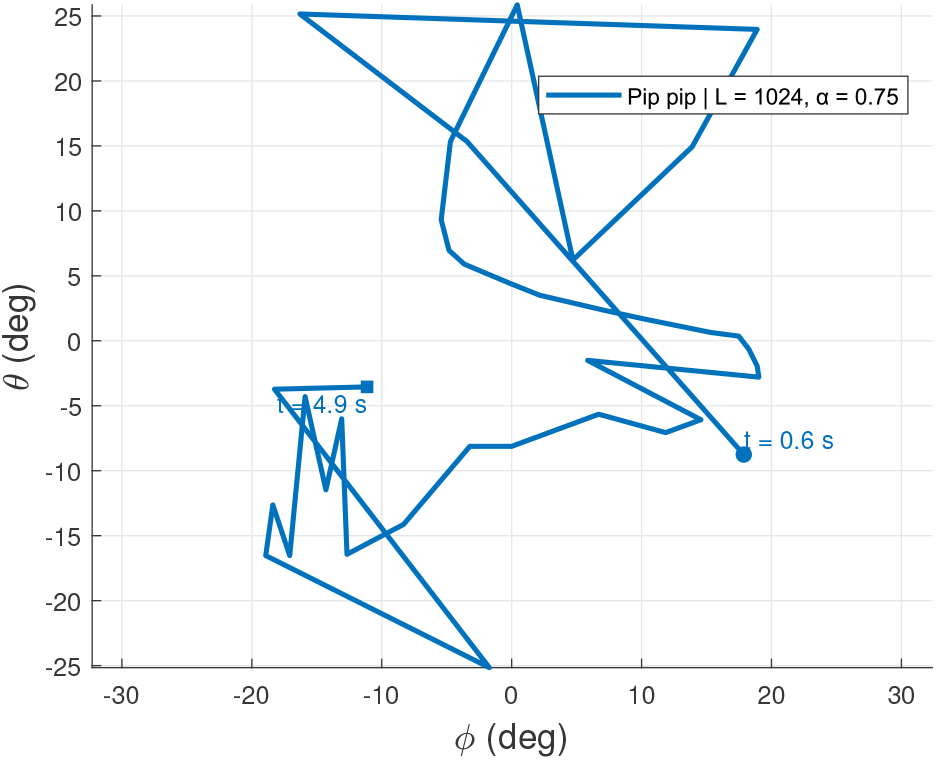
Example 2D spatial trajectory for a single recording. The trajectory corresponds to a single detected bat, where vertices occur at each moment the bat calls. Space dimensions are expressed in Polar coordinates where horizontal and vertical axes represent azimuthal and elevation angles, respectively.

To directly assess the effect of the selected configuration, Fig. 6(a) compares trajectory estimates obtained using a large analysis window (*L* = 4096, no overlap) and the proposed configuration (*L* = 1024, 75% overlap, strict frame inclusion). Both configurations recover a similar overall trajectory, but differ in their local structure and in the timing of directional changes. This demonstrates that the localisation output is sensitive to the temporal sampling configuration and highlights the need to evaluate parameter choices in terms of consistency and physical plausibility rather than visual appearance alone.

**Figure 6:**
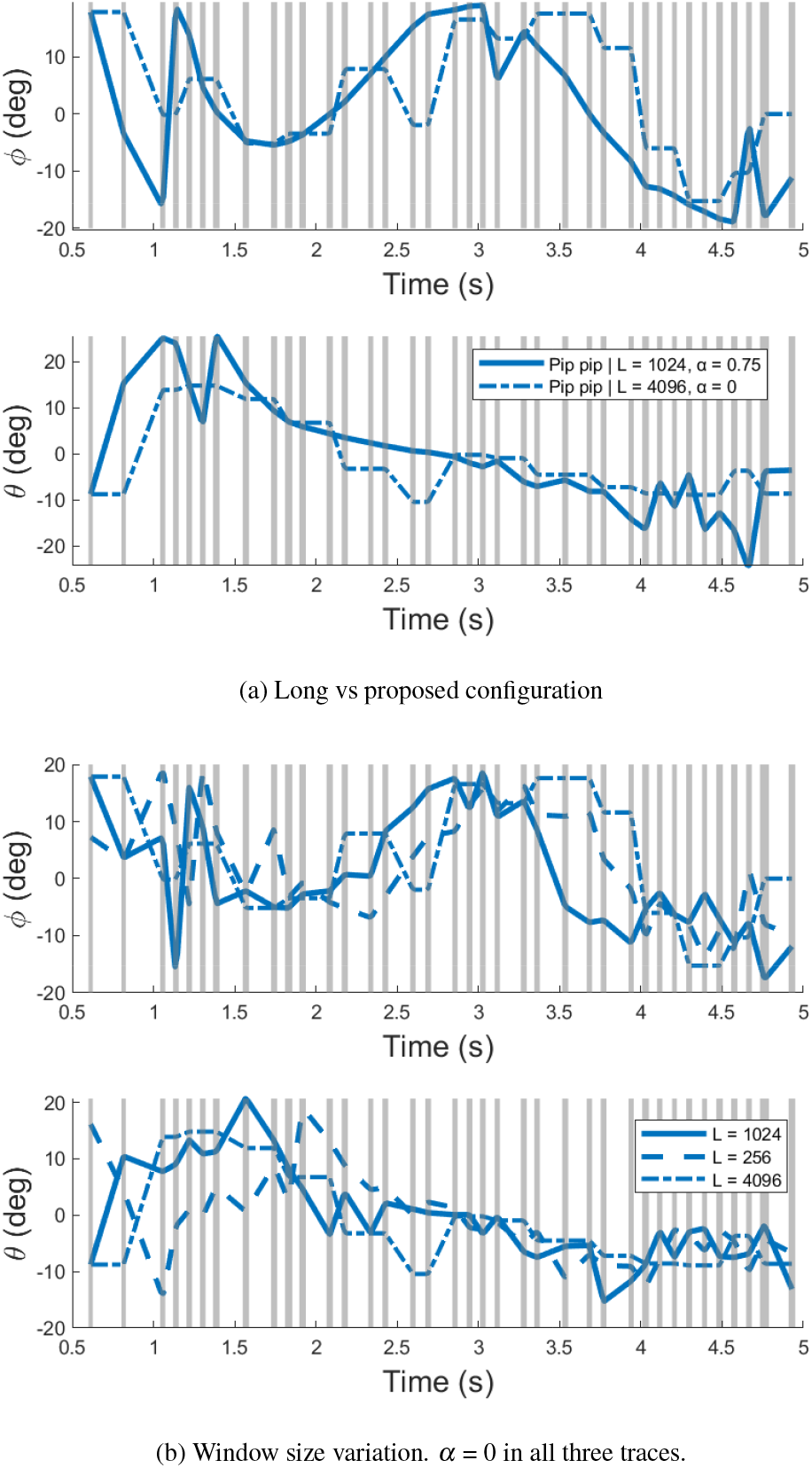
Comparison of trajectory estimates under different temporal sampling configurations: (a) comparison between a long window (*L* = 4096, no overlap) and the proposed configuration (*L* = 1024, 75% overlap, strict frame inclusion); (b) comparison between different window sizes *L* = 256, 1024, 4096 without overlap. Grey vertical bands correspond to bat calls; the width of the bands is ×5 the original one for visibility. Line style denotes the parameter configuration.

**Figure 7:**
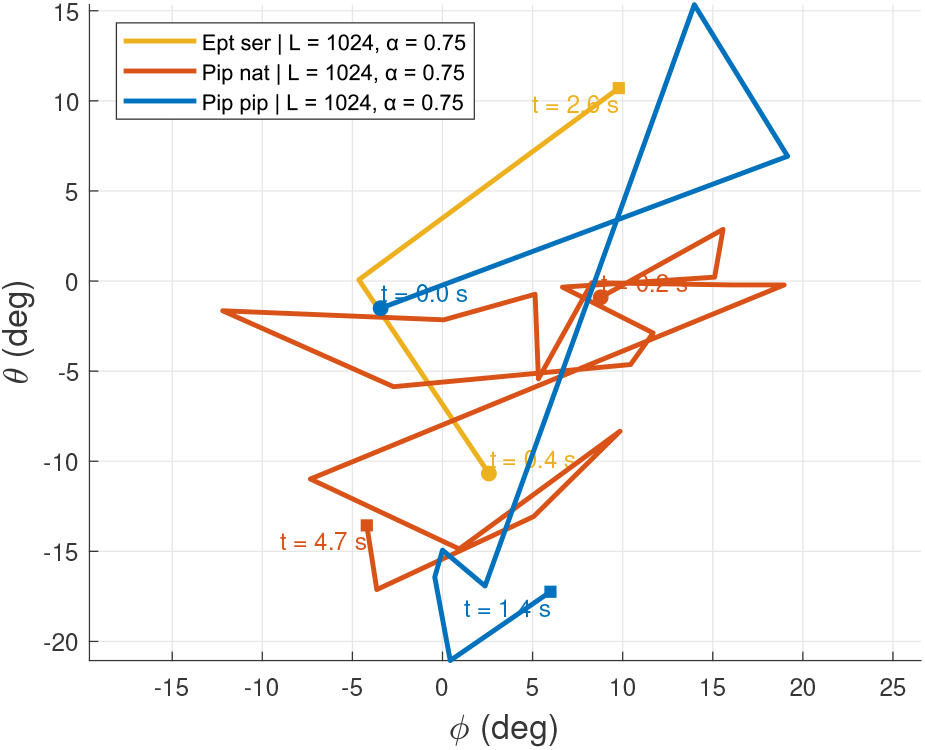
Example of reconstructed spatial trajectories for multiple simultaneously active sources within a single recording. Distinct tracks correspond to different detected events. Space dimensions are expressed in Polar coordinates where horizontal and vertical axes represent azimuthal and elevation angles, respectively.

Figure 6(b) extends this comparison across multiple window lengths. The shortest window (*L* = 256) exhibits the largest variability, whereas the longest window (*L* = 4096) produces the most stable estimates. The selected configuration (*L* = 1024) provides an intermediate behaviour, retaining local trajectory variations while avoiding the pronounced fluctuations observed at shorter window lengths.

When the estimated trajectory approaches the boundaries of the field of view, both coordinates exhibit increased variability. This behaviour is expected, as the source may lie outside the observable region and the beamformer returns maxima associated with sidelobes or aliasing artefacts rather than the true source position.

Taken together, this empirical analysis shows that trajectory quality cannot be assessed from visual smoothness alone. Instead, consistency across parameter configurations provides a more reliable indicator of localisation performance. Among the evaluated configurations, *L* = 1024 with 75% overlap and strict frame inclusion provides the most robust compromise between temporal resolution and signal support while preserving local trajectory variations.

In addition to single-source trajectories, the proposed framework can be applied to recordings containing multiple simultaneously active sources. Fig. 7 illustrates an example in which multiple spatial trajectories are reconstructed within the same recording. This demonstrates that the proposed framework remains applicable in more complex acoustic scenes with overlapping sources.

## 4. Results

### 4.1 Evaluation metrics

The performance of the proposed framework is evaluated in terms of computational efficiency and its ability to produce stable and interpretable spatial trajectories from field recordings of free-flying bats. All computations were performed in MATLAB on a 64-bit system equipped with an Intel Core i7-13700H CPU (2.40 GHz) and 32 GB RAM, and runtime measurements were obtained under these conditions.

Results are reported over a dataset of 115 recordings of 5 s each. Quantities in Table 2 are computed per recording and then averaged across the dataset, while the number of evaluated time frames, of *L* = 1024 samples each, is reported over the full dataset.

**Table 2:** Computational cost comparison between baseline continuous beamforming and BatDetect2-guided beamforming. Values are reported per recording and averaged over a dataset of 115 recordings of 5 s each; the only exception are the time frames evaluated, which refer to the full dataset. The number of time frames evaluated is reported over the full dataset.

| Metric | Baseline | Guided | Reduction |
| --- | --- | --- | --- |
| Time frames evaluated | 5.3e5 | 2.2e4 | $24 \times$ |
| Frequency bins per frame | 512 | 154 | $3 \times$ |
| Spatial grid points | 1728 | 1728 | – |
| Total evaluations | 4.7e11 | 6.2e9 | $77 \times$ |
| Wall-clock runtime (s) | 300 | 1.1 | $273 \times$ |
| Real Time Factor | 60 | 0.22 | $273 \times$ |

In the guided configuration, frequency bins per frame were determined by identifying the frequency range of detected calls in each recording and averaging the corresponding bin counts. Time frames were included only if both the start and end of the frame lay within the temporal extent of a detected call.

Spatial grid points were defined over a field of view of ±20° in azimuth and ±26° in elevation. For the runtime breakdown analysis, a grid of 36 ×48 points was used. This configuration is coarser than the dense spatial sampling discussed in Sec. 2.3, and was adopted to make the computational cost of the breakdown analysis tractable as the number of sampling points scales linearly.

Runtime measurements are reported per recording. The realtime factor (RTF) is defined as the ratio between processing time and recording duration, such that RTF = 1 corresponds to real-time operation.

### 4.2 Computational efficiency

The computational cost of high-resolution ultrasonic beamforming is dominated by the number of beamformer evaluations, which scales with the joint time-frequency-space dimensionality (Sec. 2.4). Detection-guided sparsity reduces this dimensionality by restricting processing to time-frequency regions containing bat calls.

Relative to baseline continuous beamforming, the number of evaluated time frames is reduced from approximately 5.3 ×10^5^ to 2.2 ×10^4^, reflecting an effective temporal fraction *ρ*_*t*_ = 0.0415. The number of frequency bins per frame decreases from 512 to 154, corresponding to a spectral fraction *ρ*_*f*_ = 0.3 and reflecting the band-limited nature of detected calls. The spatial grid resolution remains unchanged.

These reductions combine multiplicatively, yielding a decrease in total beamforming evaluations from approximately 6.7 ×10^12^ to 8.8 ×10^10^, corresponding to a reduction factor of *ρ*_*t*_*ρ*_*f*_ = 0.0131, consistent with the scaling expected in Sec. 2.4. Runtime decreases from 300 s per recording to 1.1 s, corresponding to a reduction factor of 0.0037. The corresponding RTF decreases from 60 to 0.22, indicating that guided beamforming operates faster than real time under the tested conditions.

Figure 8 shows that detection-guided beamforming reduces the computational burden by 2 orders of magnitude. The majority of computational cost saved is in the beamforming stage, with the cost arising from BatDetect2 comparatively negligible in the processing pipeline.

**Figure 8:**
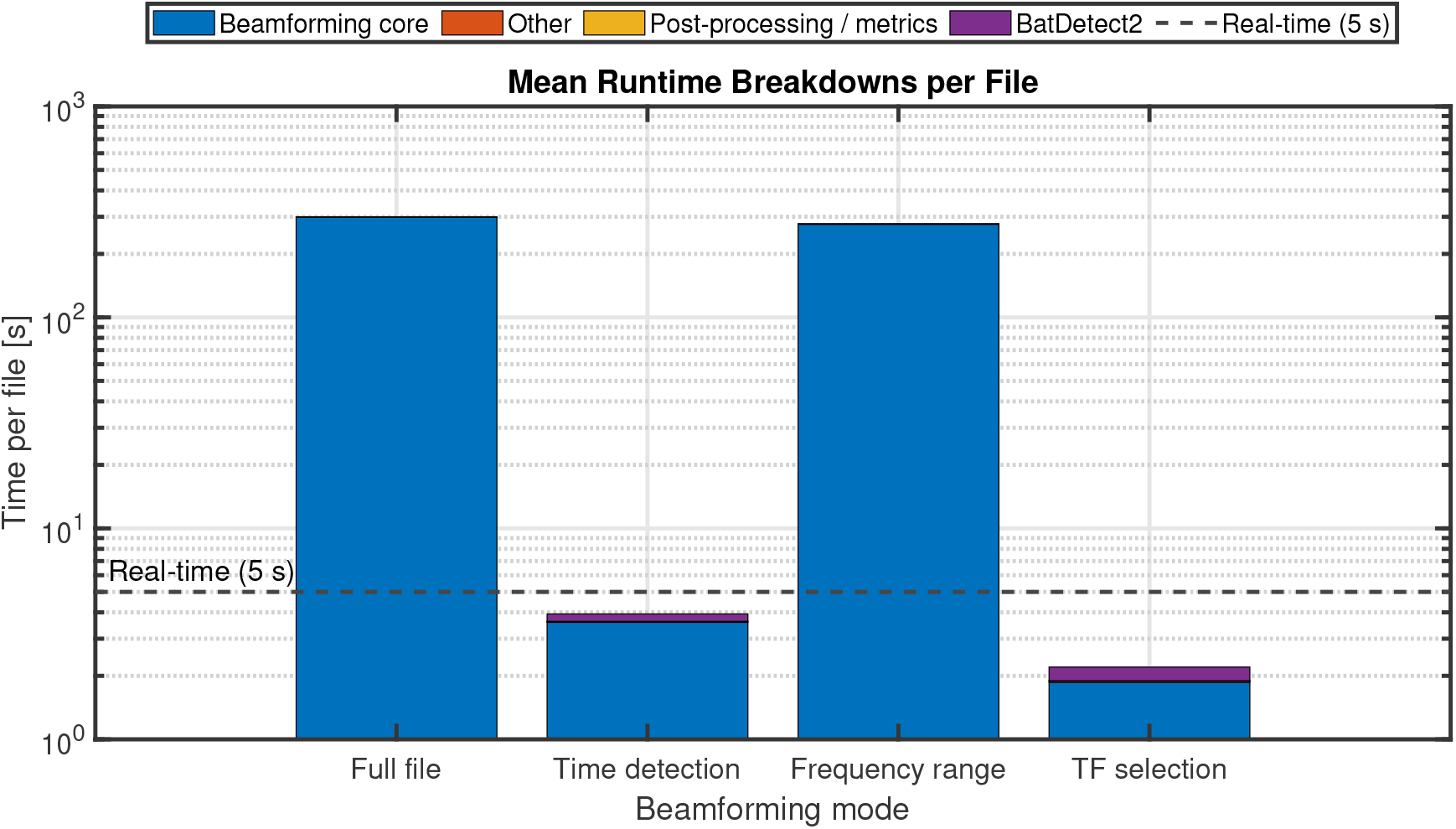
Runtime Breakdown for processing a 5 s long file.

## 5. Discussion

From an ecological perspective, the proposed framework supports a transition from activity-based monitoring, in which animal presence is inferred primarily from acoustic activity, towards spatially resolved analysis based on source location and movement. This aligns with broader developments in ecoacoustics that aim to move beyond descriptive assessments towards process-based understanding of ecological systems [1, 2]. Access to spatial trajectories enables the investigation of habitat use, flight behaviour, species interactions, and ultimately improved abundance estimation, addressing several of the applications identified for acoustic localisation systems [12]. Such information remains inaccessible with conventional single-channel approaches and supports the use of acoustic cameras for routine ecological monitoring [7, 11]. In this context, the proposed framework makes acoustic cameras more suitable for applications requiring continuous and non-intrusive observation of animal movement.

A key result of this work is the substantial reduction in the computational cost of high-resolution ultrasonic beamforming without compromising spatial resolution. This directly addresses a major limitation of acoustic localisation systems, namely the cost of exhaustive time-frequency-space processing and the challenges associated with analysing large multichannel datasets [7, 11, 12]. Reducing the real-time factor from highly super-real-time to sub-real-time operation has practical benefits for field deployment. For denser grids required to fully resolve ultrasonic beampatterns, absolute runtimes would increase, and real-time operation may not always be achieved. However, the reduction in computational cost driven by detection-guided sparsity (*ρ*_*t*_*ρ*_*f*_≈ 0.0131) is independent of spatial resolution. Consequently, although the real-time factor scales with grid density, the relative advantage over exhaustive processing is preserved.

The proposed framework also treats detection as an explicit front-end for spatial processing. Detection-guided beamforming has previously been explored in acoustic imaging contexts [31], but applications to highly transient ultrasonic signals under ecological field conditions remain limited. In contrast to approaches that combine object detection with visual image processing, the present strategy relies exclusively on acoustic information. This enables localisation in conditions where visual sensing is impractical, including nocturnal monitoring, and allows the framework to be applied to systems that do not incorporate imaging hardware. This coupling between machine-learning-based detection and beamforming also provides advantages beyond localisation. As demonstrated in [11], beamforming can improve signal-to-noise ratio by emphasising sounds arriving from target directions, potentially benefiting downstream species classification. It also points towards adaptive or species-conditioned processing strategies in which time-frequency constraints are adjusted dynamically.

Several limitations remain. First, the efficiency of the pipeline depends on both the accuracy and computational cost of the detection stage; more demanding models may offset the gains obtained through sparsity. Second, the approach assumes sparse acoustic scenes. While this holds for the recordings considered here, it may not apply in dense environments, such as roosts, where increased temporal and spectral occupancy reduces sparsity and associated efficiency gains. Third, the analysis relies on far-field and free-field assumptions. While the present data contain limited reflections, more complex environments may introduce multipath artefacts. In addition, nearby sources may violate the planar-wave approximation and require more computationally demanding formulations. Finally, the quasi-stationarity assumption remains sensitive to rapid source motion or abrupt device movement, which may introduce spatial smearing.

The array configuration was not investigated in this work. The number of microphones, aperture, and inter-sensor spacing influence resolution, aliasing, and computational cost, creating trade-offs between portability and localisation performance. Determining the minimum array requirements and spacing constraints for reliable localisation, particularly under highly nonstationary conditions, remains an important direction for future research.

Taken together, these results demonstrate that exploiting the sparsity of bat echolocation signals can substantially reduce the computational burden of high-resolution beamforming while preserving localisation performance. This improves the practical applicability of acoustic cameras for passive acoustic monitoring and supports ecological analyses that require spatial information, including habitat-use studies, behavioural monitoring, and conservation applications.

## 6. Conclusion

This work investigated how the limited time–frequency occupancy of bat echolocation signals can be exploited to reduce the computational burden of acoustic localisation. Rather than performing exhaustive beamforming across all time-frequency regions, localisation was restricted to detector-identified regions likely to contain bat calls. This reduced the number of beamformer evaluations by a factor of 77 while maintaining the spatial resolution required for trajectory reconstruction.

Beyond the computational savings, the results demonstrate that the interaction between source motion, signal non-stationarity, and short-time analysis design strongly influences localisation performance. Physically consistent temporal and spatial sampling was essential for obtaining stable trajectories, highlighting that computational efficiency alone is insufficient for reliable localisation of echolocating bats.

More broadly, these findings suggest that acoustic localisation for passive acoustic monitoring can benefit from being treated as a sparse inference problem rather than an exhaustive imaging problem. Under this perspective, computational effort can be concentrated on acoustically relevant regions while preserving localisation performance. This facilitates the extraction of spatial information from ecological acoustic recordings and may support future applications involving animal movement, habitat use, and behavioural analysis.

## Acknowledgements

This project has received funding from the European Union’s Horizon Europe research and innovation programme under the Marie Skłodowska-Curie grant agreement No. 101116715 (BioacousticAI Doctoral Network).

The authors thank Juan Sebastian Cañas (University College London) and Agata Staniewicz (Bat Conservation Trust) for their assistance with data collection.

## CRediT authorship contribution statement

Roberto Alessandri: Conceptualisation, Methodology, Software, Writing - original draft.

Lia Gilmour, Jin Jack Tan, Alejandro Osses, Dan Stowell: Supervision, Validation - review, Writing - editing.

## Data availability

Data associated with this study are available upon reasonable request.

## Declaration of Generative AI and AI-assisted technologies in the writing process

During the preparation of this work, portions of the text were drafted with the assistance of Microsoft Copilot. The authors have reviewed and edited the content and take full responsibility for the validity and integrity of the work.

## Appendix A. Beamforming derivation

Frequency-domain delay-and-sum (DAS) beamforming is a widely used and physically interpretable baseline for acoustic imaging [16, 21, 22]. The acoustic signal in free-field at the *m*-th microphone can be expressed as

**Figure Appendix B.1:**
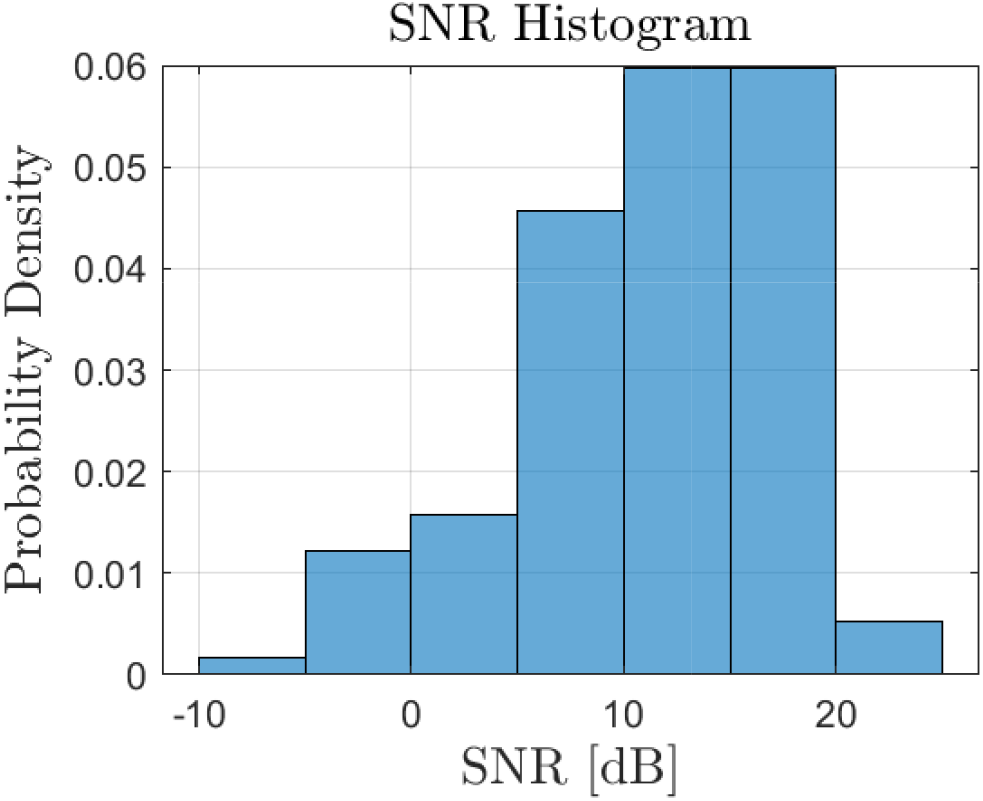
SNR Distribution of the recording session 5, used for this study

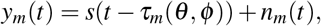

where *s*(*t*) is the source signal as a function of time *t, τ*_*m*_(*θ, φ*) is the propagation delay to the *m*-th sensor, and *n*_*m*_(*t*) denotes additive noise. Taking the Fourier transform yields

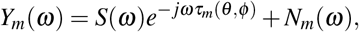

where *ω* denotes the angular frequency. Stacking all channels into the observation vector **y**(*ω*) ∈ ℂ ^*M*^, and assuming far-field propagation so that amplitude variations across the array are negligible, the steering vector is defined as

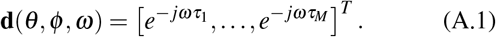

For a delay-and-sum beamformer, the weight vector is given by

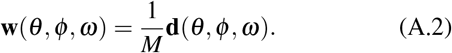

The beamformer output for a single snapshot is

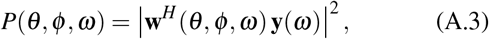

where (·)^*H*^ denotes the Hermitian transpose.

## Appendix B. Noise estimation and SNR definition

Estimating signal-to-noise ratio (SNR) for bat echolocation signals is non-trivial due to their short duration, broadband nature, and strong non-stationarity.

In this work, SNR estimation is based on a frame-level analysis. Signals are segmented into short-time frames of duration 1-5 ms, consistent with the temporal structure of bat echolocation calls. To avoid considering frequency components that were not relevant, all signals were filtered with a high-pass filter with a cutoff frequency at 10 kHz. Due to the transient and non-stationary nature of echolocation signals, noise levels are estimated using feature-based noisy-frame selection. This approach follows the general principles of signal-versus-background discrimination commonly used in probabilistic audio modelling and noise estimation [35].

The selected temporal and spectral descriptors were chosen from the audio-feature literature and include short-time energy, zero-crossing rate, kurtosis, spectral flatness, spectral entropy, spectral flux, and spectral concentration measures. Similar descriptor families are widely used for acoustic signal characterisation and are described in the Timbre Toolbox framework [36]. For each feature, candidate noisy frames are defined as those belonging to the top or bottom p-percentile of the feature distribution, depending on the expected behaviour of noise (e.g., lowest energy or highest entropy). In this work, the percentage is set in the range 15-30%.

The final set of noisy frames is defined as the intersection of the candidate sets obtained from the different features. This intersection-based approach reduces the probability of misclassifying transient signal components as noise.

Given the set of detected call frames and the set of noisy frames, SNR is estimated as:

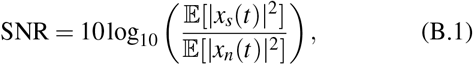

where *x*_*s*_(*t*) denotes the signal component (frames containing detected echolocation calls) and *x*_*n*_(*t*) denotes the noise component (selected noisy frames), and E[·] denotes temporal averaging.

This approach provides a data-driven estimate of SNR under field conditions without requiring explicit modelling of noise statistics and is adapted to the transient and non-stationary nature of bat echolocation signals. An example distribution of SNR distribution, corresponding to Session 5 in Tab. 1, is shown in Fig. Appendix B.1.

## Appendix C. Additional temporal sampling analysis

A critical factor when choosing the short-time analysis parameters is the frame inclusion criterion. Figure Appendix C.1(b) compares trajectory estimates obtained using strict and loose frame selection. Both approaches recover a similar overall trajectory, but differ in their local structure. Restricting processing to frames fully contained within the detected call interval preserves sharper trajectory variations, whereas looser inclusion produces a smoother estimate by incorporating frames with only partial signal content.

Figure Appendix C.1(a) illustrates the effect of overlap. In contrast to the window-length analysis, the estimated trajectories remain broadly consistent across all tested overlap values. Local discrepancies are limited to specific portions of the trace, indicating that overlap has a comparatively small influence on the localisation output within the range considered here. Increasing overlap primarily provides denser temporal sampling without substantially altering the overall trajectory.

**Figure Appendix C.1:**
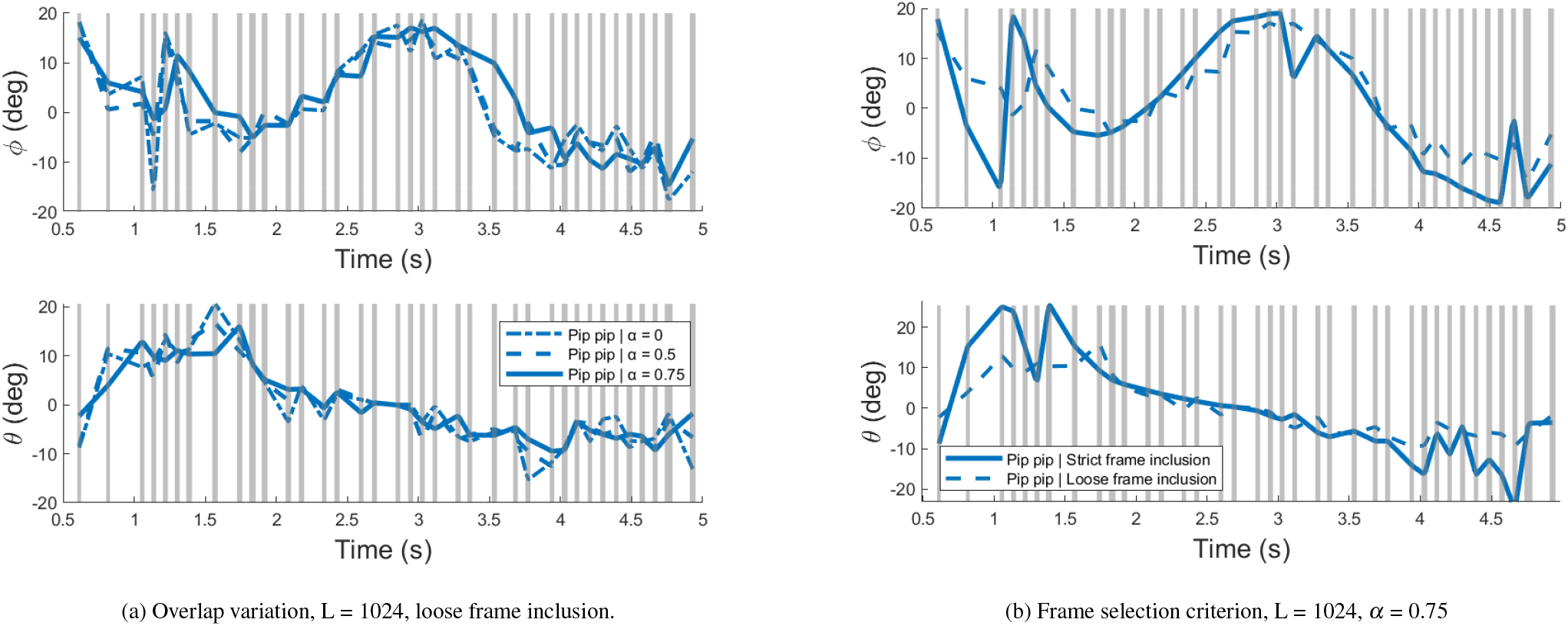
Comparison of trajectory estimates under different temporal sampling configurations: (a) comparison between different overlaps for the same *L* = 1024; (b) comparison between different frame selection criteria. Grey vertical bands correspond to bat calls; the width of the bands is 5× the original one for visibility. Line style denotes the parameter configuration.

